# Diet-derived peptides mediate the effects of dietary protein source on gastrointestinal health

**DOI:** 10.64898/2026.07.27.741049

**Authors:** Shrey D. Thaker, Lorraine Danowski, Sotiria Everett, Annemarie Ng, Xaioyue Zhang, Jie Yang, Jaclyn R. Dweck, Olga Aroniadis, Jonah S. Vadakkan, J. Alfredo Blakely-Ruiz, Ayesha Awan, Shaked Uzi-Gavrilov, Manuel Kleiner, Josephine Connolly-Schoonen, David C. Montrose

## Abstract

Plant-based diets support gastrointestinal (GI) health while animal-based diets can disrupt gut homeostasis. Although multiple aspects of these diet types are believed to confer their respective effects, the role of their protein component is less well understood. Here, we conducted a randomized crossover-controlled feeding trial wherein healthy subjects consumed 70% of their daily protein intake in the form of pea protein (PP) or egg white protein (EWP) isolate (NCT05619939). Individuals who consumed EWP reported increased GI symptoms and exhibited elevated intestinal permeability. In contrast, these endpoints did not change following PP consumption. Fecal analysis showed increased diet-derived peptides only following EWP consumption, which was associated with resistance of EWP isolate to degradation by digestive enzymes *in vitro*. Metagenomic, metaproteomic and metabolomic analyses of stool after the EWP-based diet showed reduced abundance of multiple gut-protective bacterial species and increased bacterial amino acid utilization compared to samples following the PP-based diet. Dietary peptides in the gut luminal content of EWP-fed subjects reduced metabolic function of intestinal epithelial cell in culture. Providing an amino acid-based diet mimicking EWP composition to mice prevented colonic accumulation of diet-derived proteins and GI dysfunction associated with EWP diet consumption. Collectively, these findings demonstrate that dietary protein source is a key mediator of GI function, revealing a modifiable lifestyle factor that impacts human health.

## Introduction

Disruptions in intestinal homeostasis are contributing factors to the onset and severity of gastrointestinal (GI) conditions including irritable bowel syndrome, inflammatory bowel diseases (IBD) and colorectal cancer (*1–3*). Gut luminal content, including microbiota and small molecules (host and microbially-derived), is a key mediator of GI function (*4, 5*). These factors can modify gut barrier function, modulate host-specific cell signaling pathways, and mediate immune responses (*6–9*). Diet is a major driver of microbial and metabolite profiles of the GI tract by both directly contributing diet-derived molecules and modifying bacterial abundance and/or metabolism (*10–15*). While these facts highlight the importance of the interplay between diet, gut microbes and metabolites for mediating GI health, work is still needed to elucidate the precise dietary components that mediate the balance between GI homeostasis and dysfunction.

Consumption of plant-based diets is associated with gut health while animal-based diets can impair GI homeostasis (*16, 17*). Because each of these diet types is complex, it can be challenging to discern which of their numerous components mediates their respective effects on gut health. Among the macronutrients available in diet, protein has emerged as a critical component affecting GI health. In fact, studies have shown that high protein consumption promotes gut inflammation and dysbiosis in rodents (*18–21*). Such findings have extended to humans, whereby those individuals consuming higher levels of protein – specifically derived from red meat – had increased incidence of IBD and patients with IBD experienced decreased time to relapse (*22–24*). Recent evidence in mice shows that consumption of protein isolates from beef and egg whites increases the severity of colitis, while pea-derived protein attenuates colonic inflammation (*25*). This previous work suggests that consumption of animal-based diets, especially its protein component, drives GI dysfunction. However, no prospective studies in humans have investigated this concept, or in turn, whether consumption of plant-derived protein is more beneficial for gut health.

The current study compared the impact of dietary protein derived from egg whites (EWP) *vs*. peas (PP) on the gut health of human subjects. EWP consumption increased GI symptoms and permeability while no changes in these endpoints were found following the PP feeding stage. These EWP-induced effects were associated with increased diet-derived peptides and alterations in microbial populations and metabolism. *In vitro* and *in vivo* studies demonstrated that the excess colonic accumulation of protein following EWP consumption directly impaired intestinal epithelial cell function and reduced gut health. Notably, PP consumption did not exert such adverse effects on gut health or intestinal epithelial cell metabolism. Taken together, these findings suggest that dietary protein source is a significant contributor to GI homeostasis and provide support for the importance of diet for mediating human health.

## Results

### Subjects demonstrated strong adherence to study diets

Following a rigorous screening process, 18 individuals were enrolled in this randomized controlled-crossover trial, with one subject dropping out after the first phase and 17 subjects completing the entire dietary intervention (**Fig. 1**). Subjects were given diets that were formulated to meet calorie needs and provide 1.5g of protein per kg body weight with 70% of daily protein intake derived from PP or EWP isolate blended into smoothies, while the remainder of the diet consisted of whole foods. Each protein source was given for a period of 10 days followed by a 14-day washout period after which subjects were given the other source for 10 days (**Fig. 2**). Body weight during the dietary intervention stayed within 0.5% of baseline and was similar for PP and EWP phases (-0.5 ± 0.9kg *vs*. -0.3 ± 0.7kg, respectively) indicating that the diets met subjects’ calorie needs. Total calorie and protein consumption per day was similar when subjects were in either the PP or EWP-diet phases (**Table 1**). Calories and protein consumed by subjects from food and smoothies closely met the goals of the intervention (**Table 1**). Energy and protein consumption per subject is shown in Table S1. Consumption of foods

**Fig. 1.**
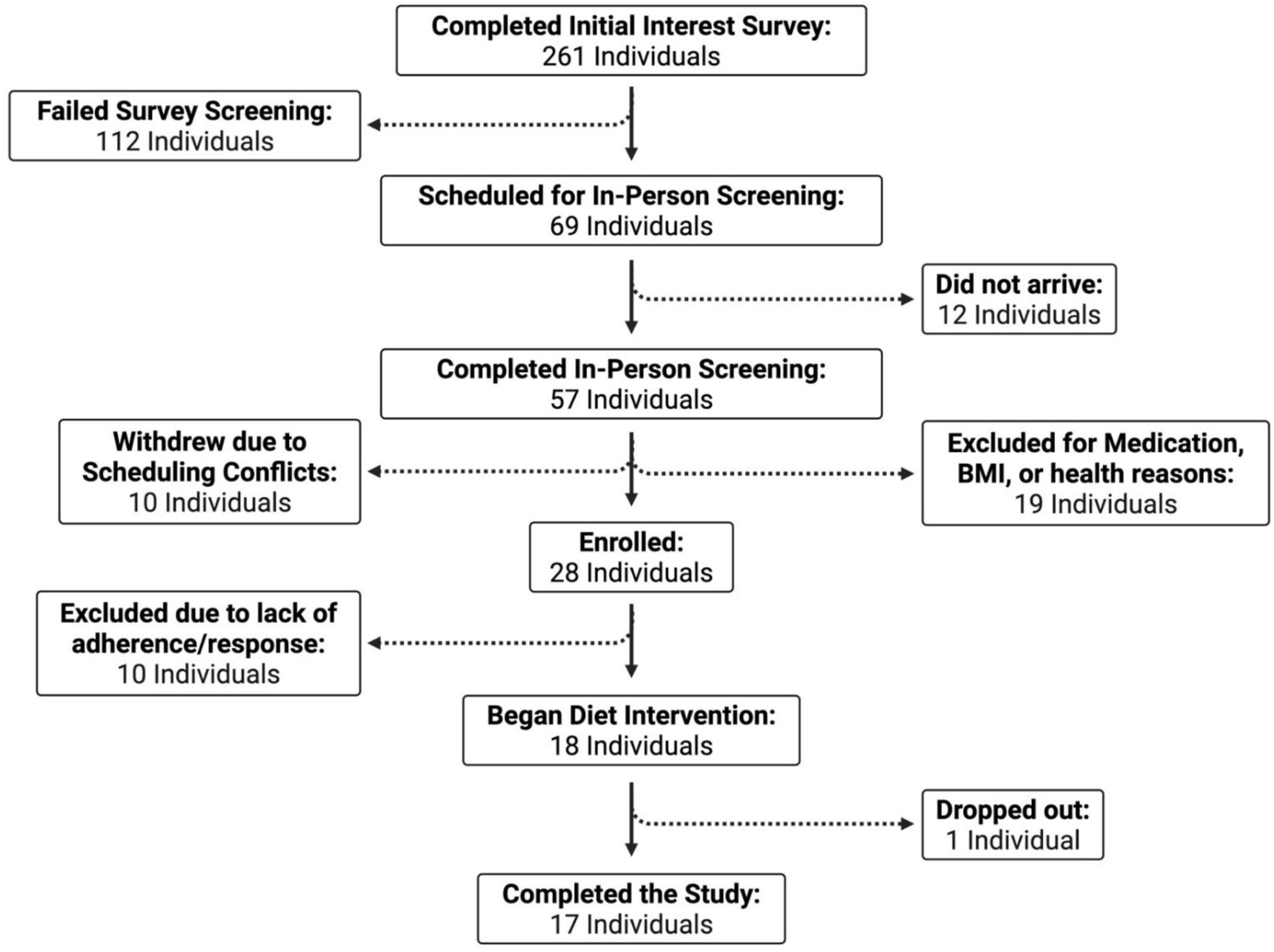
CONSORT diagram. Created in BioRender.

**Fig. 2.**
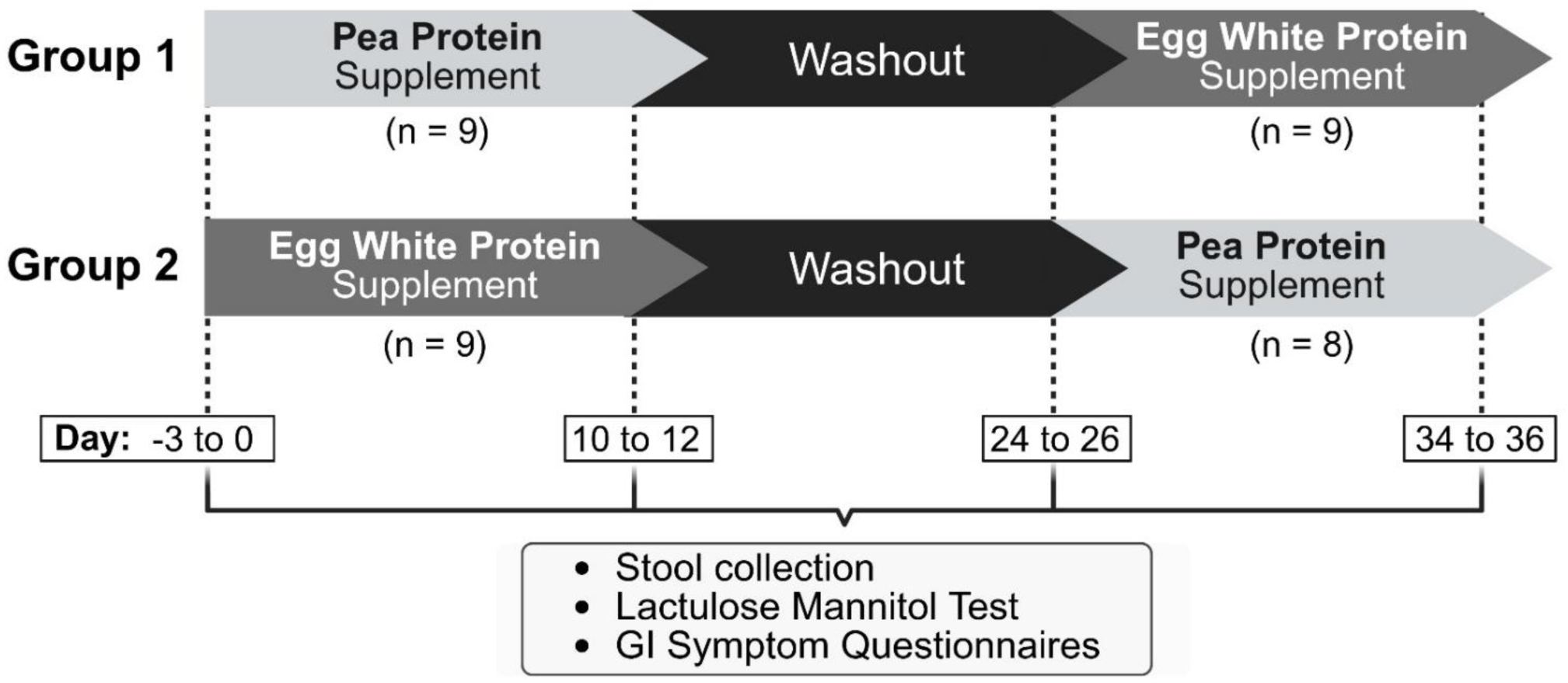
Trial design. Created in BioRender.

**Table 1.** Study diet nutrient source, intake, and adherence.

|  | **Pea Protein Phase (n=17)** |  | **Egg White Protein Phase (n=18)** |  |
| --- | --- | --- | --- | --- |
|  | Consumed | Percent of Goal | Consumed | Percent of Goal |
| **Energy** |  |  |  |  |
| From food |  |  |  |  |
| *kcal/day* | 1105.4 ± 243.3 | - | 1164.9 ± 262.1 | - |
| *kcal/kg/day* | 15.6 ± 3.8 | - | 16.6 ± 4.5 | - |
| From smoothie |  |  |  |  |
| *kcal/day* | 770.6 ± 141.5 | - | 758.7 ± 129.9 | - |
| *kcal/kg/day* | 10.1 ± 2.6 | - | 10.6 ± 0.5 | - |
| Total |  |  |  |  |
| *kcal/day* | 1876.0 ± 312.7 | 97.7 ± 10.4 | 1923.4 ± 313.6 | 99.2 ± 10.0 |
| *kcal/kg/day* | 26.3 ± 4.0 | 97.7 ± 10.4 | 27.2 ± 4.8 | 99.2 ± 10.0 |
| **Protein** |  |  |  |  |
| From food |  |  |  |  |
| *g/day* | 27.9 ± 6.5 | 85.4 ± 10.2 | 27.2 ± 4.4 | 84.4 ± 8.5 |
| *g/kg/day* | 0.4 ± 0.10 | 85.4 ± 10.2 | 0.4 ± 0.0 | 84.4 ± 8.5 |
| From Smoothie |  |  |  |  |
| *g/day* | 73.8 ± 13.4 | 97.2 ± 5.0 | 74.0 ± 12.7 | 98.0 ± 5.0 |
| *g/kg/day* | 1.02 ± 0.05 | 97.2 ± 5.0 | 1.0 ± 0.1 | 98.0 ± 5.0 |
| Total |  |  |  |  |
| *g/day* | 105.5 ± 18.9 | 97.3 ± 5.8 | 105.8 ± 17.5 | 98.2 ± 5.7 |
| *g/kg/day* | 1.46 ± 0.09 | 97.3 ± 5.8 | 1.5 ± 0.1 | 98.2 ± 5.7 |

and beverages other than those provided for the study contributed minimally to calorie and protein intake (1.3 ± 2.2% and 1.4 ± 3.4% respectively, in the PP phase and 1.4 ± 1.5% and 0.5 ± 0.7% respectively, in the EWP phase).

### Consumption of egg white protein induces intestinal symptoms and permeability

To evaluate whether either dietary protein source impacted GI health, subjects were provided PROMIS® (Patient-Reported Outcomes Measurement Information System) surveys consisting of four domains related to GI symptoms. Analysis of responses revealed that while 10-days of consuming the PP-based diet did not impact symptoms, EWP consumption increased total GI symptom score, including increases in diarrhea, belly pain and gas and bloating (**Fig. 3, A-E**). To next examine the impact of protein source on gut permeability, participants were administered a lactulose:mannitol test. While no significant change was observed following PP consumption, the lactulose:mannitol ratio significantly increased in those individuals consuming EWP, indicating increased intestinal permeability (**Fig. 3F**). These findings suggest that consuming EWP as the major dietary protein source adversely impacts gut health, while dietary PP maintains gut homeostasis.

**Fig. 3.**
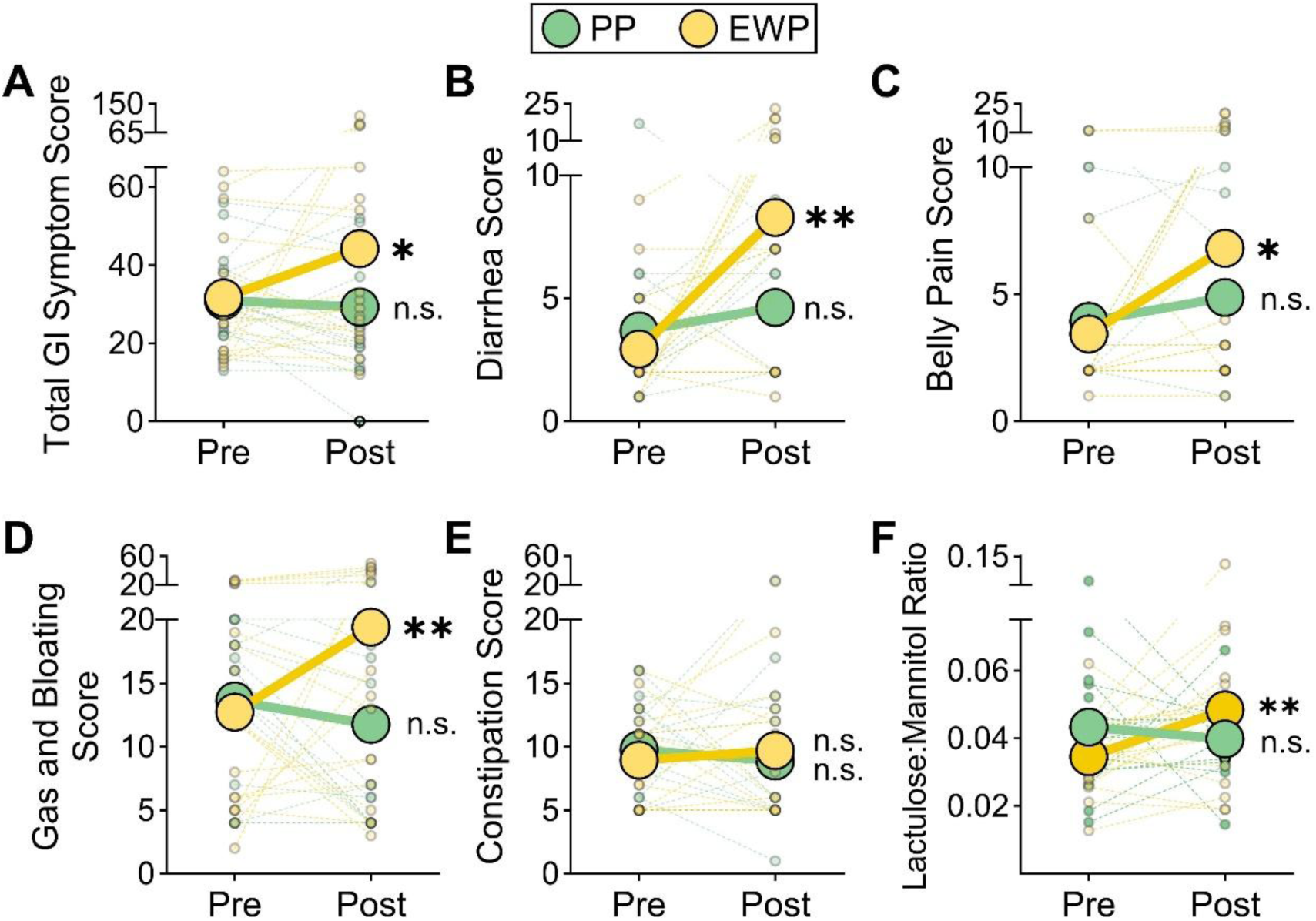
**Egg white protein consumption disrupts intestinal health while consuming pea protein maintains gut homeostasis. (A-E**) Subjects completed GI PROMIS score questionnaires before (Pre) and after (Post) a 10-day intervention consuming ∼70% of daily protein derived from isolates of egg whites (EWP) or pea (PP). Scores for total GI symptoms (A), diarrhea (B), belly pain (C), gas and bloating (D), and constipation (E) were calculated. (**F**) The ratio of lactulose:mannitol in urine following consumption of these sugars was determined Pre and Post diet intervention from subjects described in panels A-E. Dashed lines represent individual subjects and solid lines represent the average for each group. *P≤0.05; **P<0.01 comparing change in score from pre to post for each diet. n.s. = not significant.

### Egg white-derived peptides accumulate in the colonic lumen following consumption

Given that diet can shape the gut luminal environment, metabolomic profiling was carried out on feces from subjects before and after being provided EWP or PP-based diets. This analysis revealed that 10-days of feeding either protein source decreased the abundance of a similar number of metabolites, while EWP increased the levels of twice as many metabolites as PP (**Fig. 4A**). Principal component analysis (PCA) of small molecule changes showed that the EWP group exhibited a unique shift in PC2, a component largely comprised of amino acid (AA)- related molecules and peptides (**Fig. 4, B-C**). This select increase in AAs and peptides was confirmed by comparing the number of metabolites changed in abundance in these categories to the number changed within other superpathways (**Fig. 4D**). Specific molecules that comprise the “Amino Acids” and “Peptides” superpathways are shown in Figures S1 and S2. To determine the source of increased peptides following EWP consumption, metaproteomics – which discerns the relative contributions of microbial, host and diet-derived proteins in biological samples (*26*) – was employed. This analysis revealed that while no changes in the contributions of any of these sources occurred in feces following 10-days of consuming a PP-based diet, diet-derived proteins selectively increased after EWP consumption (**Fig. 4, E and F**). Moreover, among all diet-derived proteins identified in feces, over 50% mapped specifically to those proteins found in eggs following the EWP diet, compared to only 10% that mapped to proteins from peas in the PP-diet group (**Fig. 4G**). An *in vitro* system was then employed to determine whether the observed higher levels of colonic diet-derived peptides following EWP consumption, relative to PP, result from differential susceptibility of the isolate to digestive proteases in the upper GI tract. Incubation of either isolate with pepsin (gastric protease) resulted in nearly equivalent degradation (indicated by decreases in large peptides and accumulation of small peptides), while subsequent exposure to trypsin (major small intestinal protease) showed rapid degradation of PP but no further degradation of EWP isolate (**Fig. 4, H and I**). Incubation with only trypsin recapitulated the observed difference in protein degradation (**Fig. S3**). Collectively, these data suggest that compared to pea-derived proteins, proteins from egg whites are resistant to digestive enzymes resulting in impaired absorption in the small bowel, and in turn, are deposited into the colon to a greater extent.

**Fig. 4.**
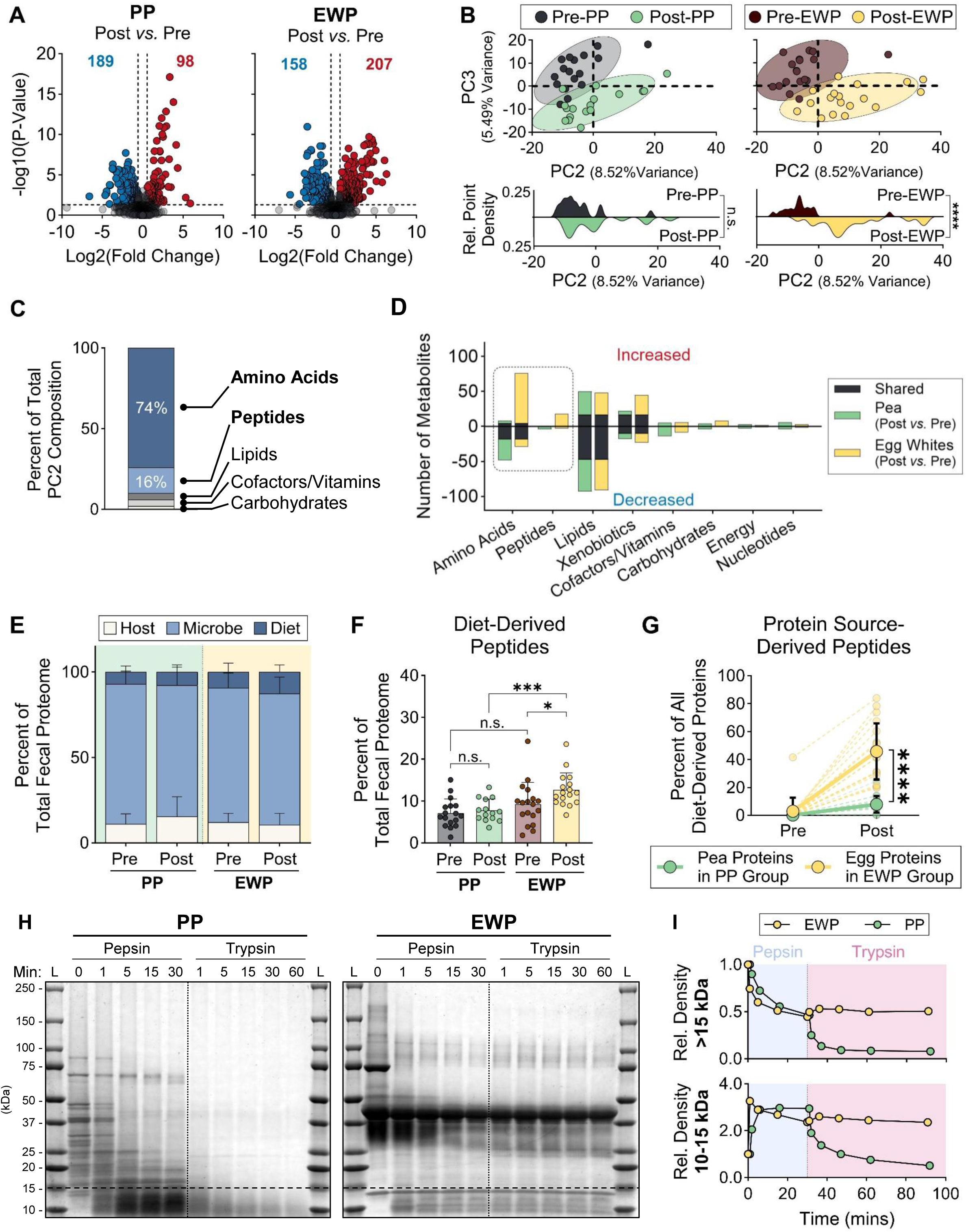
**Increased diet-derived peptides in colons from subjects consuming egg white protein. (A-D**) Metabolite profiling was carried out on feces from individuals before and after consuming diets in which the majority of protein content was comprised of isolates from egg whites (EWP) or pea (PP). (A) The significantly increased or decreased metabolites following PP (left side) or EWP (right side) consumption are shown. Blue color indicates decreased metabolites, while red color indicates increased metabolites. (B) Principal component analysis of fecal metabolites following PP (left side) or EWP (right side) consumption is shown. Histograms (bottom) show PC2 distribution of each group. (C) The superpathways represented by the top 100 metabolites within PC2 are shown. (D) Increased or decreased metabolites were categorized by superpathway and shown as uniquely decreased or increased, or shared, following either diet intervention. (**E**) The contributions of host, microbe, and diet-derived peptides relative to the total fecal proteome before and after the diet intervention was determined using metaproteomics. (**F**) Data shown in panel E were used to selectively display the relative contributions of diet-derived peptides to the total fecal proteome before and after the diet intervention. (**G**) The abundance of peptides that map to pea proteins (green) or egg proteins (yellow) as a percentage of all diet-derived proteins in the feces, was determined by metaproteomics and shown before (Pre) and after (Post) consuming the PP or EWP diets, respectively. Dashed lines represent individual subjects and solid lines represent the average for each group. (**H-I**) PP and EWP isolates were incubated with pepsin then trypsin and protein bands were visualized over time on an SDS-PAGE gel stained with Coomassie Blue G-250 (H) and the relative density of bands above or below 15 kDa was quantified (I). *P≤0.05; ***P<0.001; ****P<0.0001. n.s. = not significant.

### Consuming an egg white protein-based diet is associated with distinct gut bacterial populations and increased bacterial protein utilization

Given the differences in luminal protein abundance driven by the two dietary protein sources, we next evaluated the impact of these changes on the microbiome. Initially, metagenomics was used to measure the taxonomic abundance of the microbiota. At the phylum level, lower abundance of *Firmicutes* and higher abundance of *Bacteroidetes* were observed in the EWP group compared to the PP group, resulting in a lower Firmicutes:Bacteroidetes ratio (**Fig. 5, A and B**). Measurements at the species level revealed decreased *Eubacterium rectale*, *Roseburia inulinivorans* and *Roseburia intestinalis,* among others, in the feces of EWP-fed subjects compared to feces from individuals fed PP (**Fig. 5C**). No significant differences in overall bacterial diversity were observed between groups (**Fig. S4**). Next, we determined whether the higher abundance of diet-derived peptides in the colons of EWP-fed subjects impacted the metabolism of bacteria. Metagenomics and metaproteomics, to measure bacterial gene and protein abundance, respectively, revealed that multiple pathways related to AA biosynthesis were lower in abundance in the EWP group compared to the PP group (**Fig. 5, D and E**). Conversely, genes and proteins that drive AA degradation were significantly higher in abundance following EWP consumption *vs.* PP consumption (**Fig. 5, F and G**). Moreover, the products of bacterial AA degradation (*i.e.*, putrefaction) were elevated in the feces of the EWP *vs.* PP group, suggesting that bacteria are utilizing the abundant AAs (**Fig. 5H**). Collectively, these findings suggest that the higher levels of diet-derived peptides following EWP consumption drive distinct microbial changes.

**Fig. 5.**
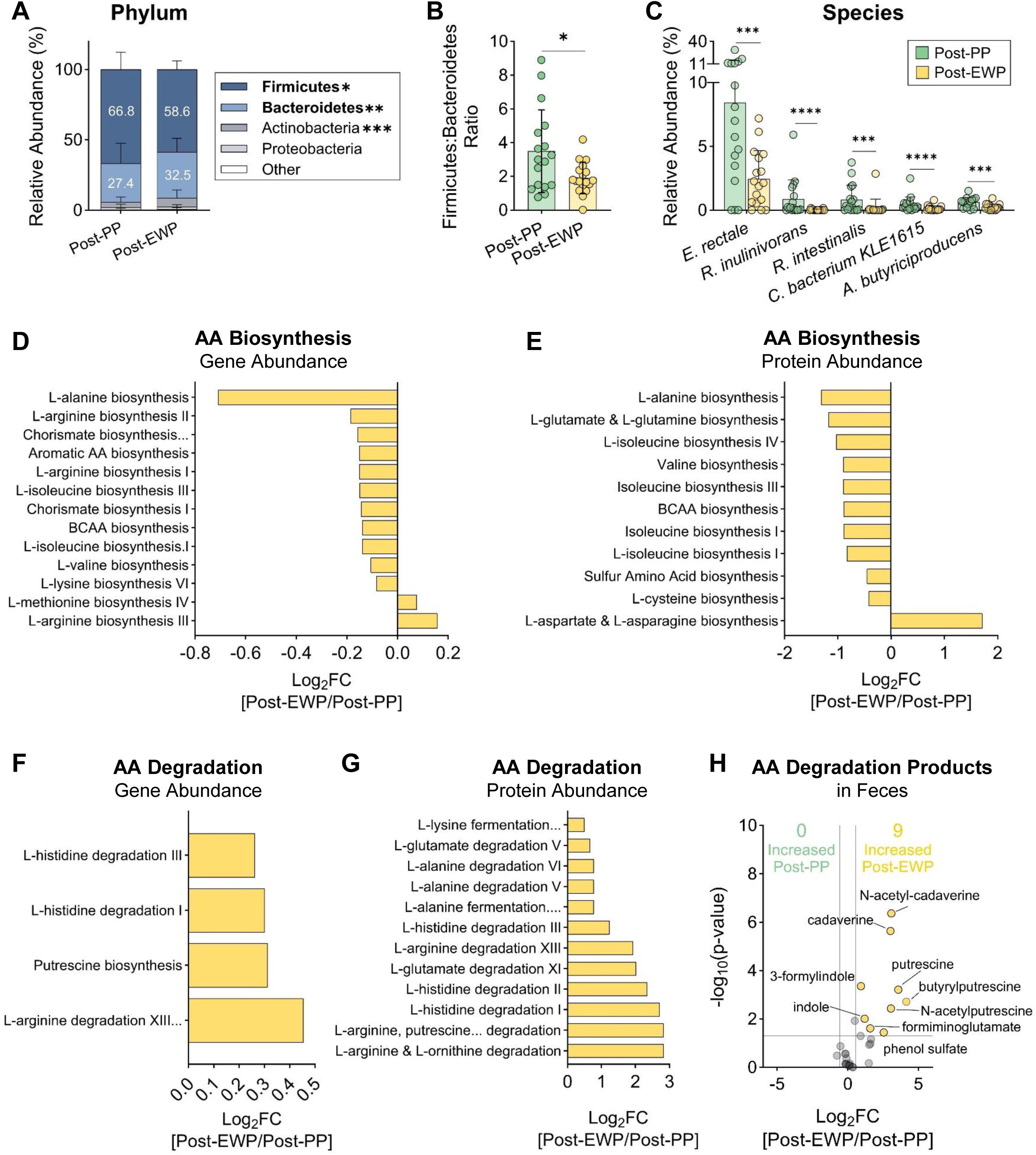
**Bacterial populations and metabolism respond to increased diet-derived peptides. (A-C**) Metagenomics was carried out on feces from subjects after 10 days of consuming egg white protein (post-EWP)- or pea protein (post-PP)-containing diets to determine the relative abundance of bacteria by phylum (A), Firmicutes:Bacteroidetes ratio (B) and species (C). (**D-E**) Log2 fold-change (FC) of pathways associated with amino acid (AA) biosynthesis in bacteria based on gene (D) and protein (E) abundance comparing post-EWP to post-PP, is shown. (**F-G**) Log2 FC of pathways associated with AA degradation in bacteria based on gene (F) and protein (G) abundance comparing post-EWP to post-PP, is shown. Values below zero represent a decrease while values above zero represent an increase. (**H**) Log2 FC of the levels of bacterial putrefaction by-products as measured by metabolomics comparing post-EWP to post-PP, is shown. *P≤0.05; ***P<0.001; ****P<0.0001.

### Gut luminal peptides derived from dietary egg white protein impair intestinal epithelial cell metabolism and sensitize mice to colonic injury

The data described above show that EWP consumption alters the gut luminal environment, including increased diet-derived peptides and changes in microbial populations and their metabolites. To determine whether these changes are directly capable of impairing intestinal epithelial cell function (a key regulator of gut health), we generated clarified, sterile-filtered supernatants from homogenized feces (hereafter called ‘fecal extract (FE)’) from subjects following 10 days of EWP or PP consumption. Applying this material to intestinal epithelial cells in culture mimics exposure of the soluble component of the gut luminal environment to the intestinal epithelium, while the MTT assay informs on the metabolic activity and health of cells. This analysis showed that, compared to FE from samples of PP-fed subjects, FE from EWP-fed individuals significantly decreased MTT absorbance (**Fig. 6A**). Next, to determine the relative contributions of peptides *vs*. small molecules for exerting this effect, EWP FE was passed through a size exclusion filter to separate molecules smaller than 3 kDa (primarily small molecules) or larger than 3 kDa (primarily peptides). This revealed that the fraction predominantly containing peptides decreased MTT to a level equivalent to the total extract, while the small molecule fraction had no effect (**Fig. 6, B and C**). To additionally assess whether diet-derived peptides mediate EWP-driven intestinal epithelial cell and GI dysfunction, we formulated an AA-based diet for mice that mimics the AA composition of EWP but does not contain whole proteins from EWP isolate (**Table S2**). This diet, which was employed to reverse dietary peptide accumulation in the colon, was fed to mice along with custom purified diets containing their entire protein content from either whole PP or EWP isolate for one week (**Table S2** and (*25*)). Diet consumption and body weight were comparable across groups during the feeding period (**Fig. S5**). First, we demonstrated that EWP diet feeding resulted in higher fecal protein levels compared to PP feeding (**Fig. 6D**). However, providing the AA-based version of EWP prevented colonic luminal protein accumulation (**Fig. 6D**). Further, we demonstrated that the observed differences in protein abundance were driven by diet-derived peptides, rather than those from microbe or host, similar to what was observed in humans (**Fig. 6, E and F**). We next applied FEs from mice fed each of these diets to intestinal epithelial cells and carried out MTT assay after confirming their protein levels paralleled those found in the respective fecal samples (**Fig. S6**). Here, we found that FE from mice fed EWP exerted a markedly greater impairment in intestinal epithelial cell metabolic function compared to FE from the PP group, while this effect was reversed with FE from mice fed the AA-based EWP diet (**Fig. 6G**). Finally, we determined whether the observed reversal of intestinal epithelial cell impairment when the deposition of EWP into the colon is prevented would extend to the *in vivo* setting. To test this, mice were fed the three diets described above for one week then challenged with the colonic epithelial irritant dextran sodium sulfate, while continuing their respective diets. Here, we found that while EWP feeding led to higher diarrhea and rectal bleeding scores compared to PP feeding, these features in mice fed the AA-based EWP diet were equivalent to PP-fed mice (**Fig. 6, H and I**). Altogether, these findings suggest that excess colonic levels of diet-derived protein, mediated by dietary protein source, adversely affect intestinal epithelial cell function and gut health.

**Fig. 6.**
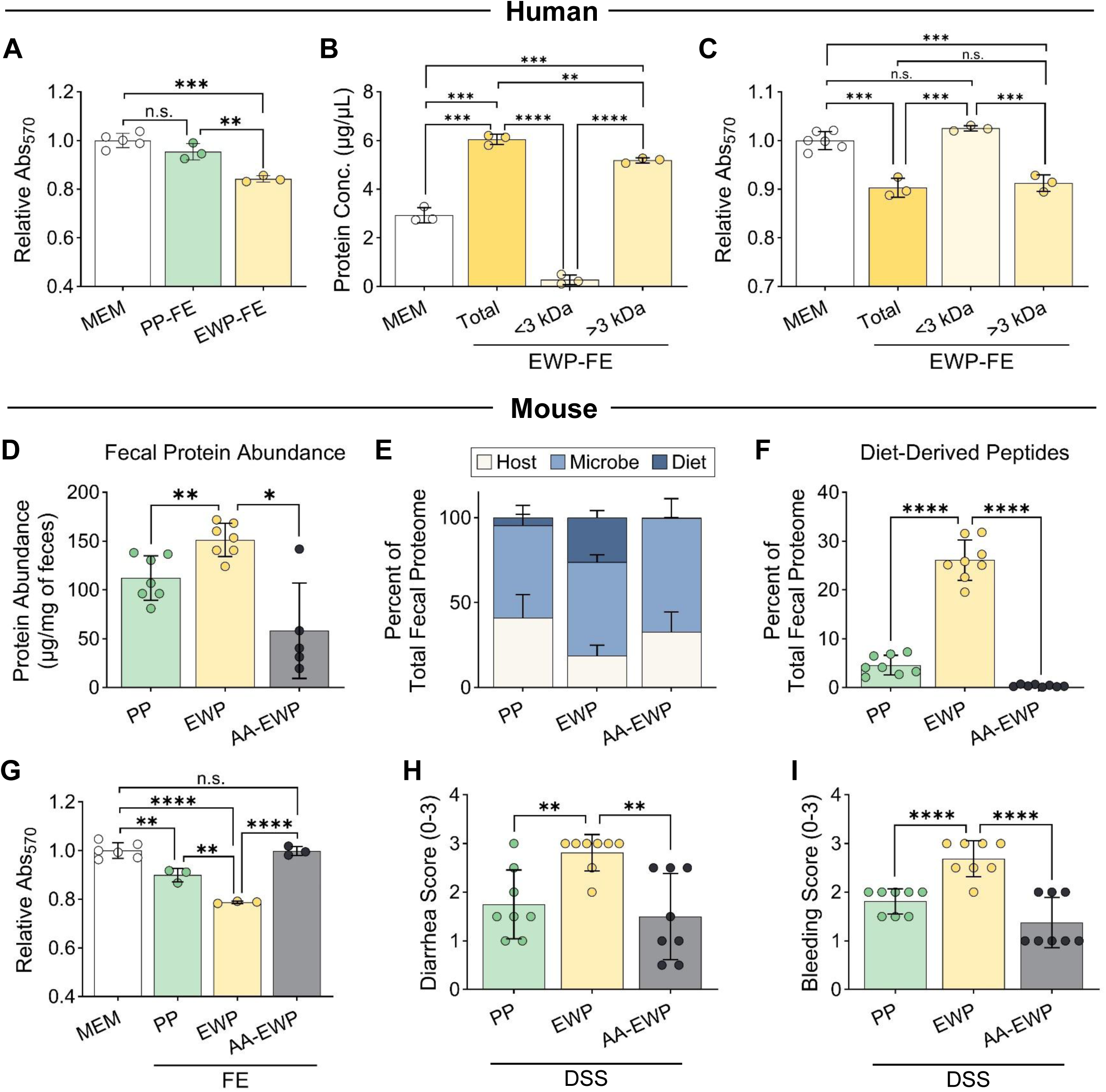
Accumulation of dietary peptides from egg white protein impairs intestinal epithelial cell metabolism and resistance to colonic injury. **(A**) IEC-6 intestinal epithelial cells were treated for 24 hours with sterile extracts from fecal samples (FE) of human subjects after consuming pea protein (PP) or egg white protein (EWP)-based diets and relative absorbance at 570 nm compared to untreated cells (MEM) was measured following MTT assay. (**B-C**) Fecal extracts from subjects after consuming EWP-based diet (EWP-FE) were filtered by size (above or below 3 kDa). Protein levels were measured in each fraction (B) and each fraction was applied to IEC-6 cells followed by MTT assay, as described in panel A (C). (**D-F**) C57BL/6J mice (n = 7-8 per group) were fed a semi-synthetic diet containing either EWP isolate, PP isolate, or an amino acid-based diet in the relative proportions found in EWP isolate (AA-EWP) for one week and fecal samples were collected. Protein abundance was measured in feces (D), contributions of host, microbe, and diet-derived peptides relative to the total fecal proteome was determined by metaproteomics (E), and the data in panel E are shown as the relative contribution of diet-derived peptides to the total fecal proteome is shown (F). (**G**) FEs were generated from samples collected from mice in the study described in Panel D and applied to IEC-6 cells to perform MTT assay, as described in Panel A. (**H-I**) Mice were fed the diets described in panel D for 1 week then given 1% dextran sodium sulfate in drinking water while continuing on their respective diets. Average diarrhea (H) and bleeding (I) scores were calculated after 5 days. *P<0.05; **P<0.01; ***P<0.001; ****P<0.0001; n.s. = not significant.

## Discussion

Plant-based diets are more beneficial for gut health than animal-based diets, although it is unclear whether the protein components of these diet styles play a role. Here, we provide evidence from a randomized controlled-feeding trial in healthy subjects that consumption of EWP induces GI symptoms and intestinal permeability whereas consuming PP maintains gut homeostasis. EWP feeding additionally increases diet-derived AAs and peptide levels in the colon, which was associated with select alterations in microbial populations and metabolism.

Moreover, EWP-driven gut luminal changes reduced intestinal epithelial cell metabolism, while limiting colonic deposition of EWP-derived dietary peptides reversed this effect, as well as gut dysfunction in mice.

Dietary protein is predominantly digested in the small intestines, where it is subsequently absorbed, thus limiting the abundance of peptides and AAs in the colonic lumen (*27*). Our data suggest that impaired digestion, and in turn, limited absorption of EWP isolate drives the observed increase in diet-derived peptides found in feces following EWP diet consumption. This concept is supported by the fact that EWP isolate exhibits impaired breakdown when directly incubated in the GI protease trypsin. Inferior digestibility of EWP compared to PP using a similar *in vitro* model system has been reported previously (*28, 29*), along with studies finding differential levels of diet-derived peptides in feces of mice depending on the protein source incorporated into diet (*26*), suggesting that differences in digestion carry over to the *in vivo* setting. The current study now extends this phenomenon to humans for the first time. Although it is not entirely clear what characteristics of EWP make it more resistant to digestion, it is possible that proteins in egg whites contain regions resistant to trypsin-mediated cleavage.

Differences in post-translational modifications may also play a role. Certainly, in-depth characterization of the structure and composition of dietary protein sources will shed light on the observed differences. Moreover, expanding analysis of digestibility to additional protein sources will help to inform on which may be favorable for maintaining gut health and whether this characteristic goes beyond simply categorizing them by animal *vs*. plant origin.

The findings shown here and previous studies demonstrate that spillover of dietary components normally absorbed in the upper GI tract into the colon alters the gut luminal environment (*19, 26, 30, 31*). In turn, the resident microbiota and host colonic mucosa are exposed to an altered molecular profile. The observed differences in bacterial populations comparing individuals after consuming EWP or PP highlight the impact that dietary peptide abundance can have on the microbial milieu of the gut. Moreover, bacterial metabolism related to AAs was profoundly affected, with the higher abundance of dietary peptides resulting in a simultaneous decrease in AA synthesis and increase in AA degradation. These findings suggest that those microbes that cannot tolerate a high protein environment are diminished in abundance while those that remain utilize peptides and AAs as a fuel source. Because those subjects consuming EWP exhibited lower abundance of bacteria shown to preserve gut health and protect against inflammation (*32–34*) compared to those consuming a PP diet, and bacterial AA degradation products can be proinflammatory (*35–38*), it stands to reason that diet-induced alterations in microbiota are playing a role in the observed phenotypes. In fact, the ability of macronutrient modulation to drive microbially-mediated alterations in GI inflammation was demonstrated in mice with high fructose feeding (thereby elevating colonic luminal fructose levels) and more recently, upon altering protein source (*25, 31, 39*). Conversely, consumption of PP may help to maintain the abundance of gut-protective species (*e.g., E. rectale*, *Roseburia* species), a concept supported by studies in a model system exposing pea proteins directly to human derived-microbes (*40*).

In addition to these bacteria-related phenomena, *in vitro* experiments conducted here suggest that direct exposure of dietary peptides to intestinal epithelial cells can impair their metabolism. In fact, removal of dietary peptides from fecal extract either directly through size exclusion or feeding an AA-based diet prevented these detrimental effects. Notably, the ability of diet-derived peptides to negatively impact the morphology and metabolism of colonocytes was shown during high protein diet feeding in rats (*41*). Although it is not clear what the precise mechanism is by which dietary peptides adversely affect intestinal epithelial cells, it is possible that these molecules are taken into cells *via* physiological endocytosis and, when present at high levels, saturate the cells’ capacity to modify or catabolize them, resulting in cell stress. In theory, the resistance of EWP proteins to physiologic digestion may extend to digestion within cells, thus perpetuating exogenous peptide accumulation. Careful interrogation of dietary peptide transport and intracellular consequences of their uptake may inform on the precise mechanisms underlying these effects. Regardless, given our findings and the diet-mediated changes in bacteria discussed above, it is possible that the observed phenotypes are driven by both host-intrinsic and microbially-mediated mechanisms.

Previous studies suggest that plant-based diets promote gut health, effects that are believed to be driven by the increased fiber and polyphenols as well as the lower saturated fat and animal-derived protein content of this diet type (*42–44*). Our findings provide the first evidence that their protein component, independent of other macronutrients, may play a role in promoting gut homeostasis in humans. This conclusion stems from the fact that PP and EWP diets were matched for macronutrient content, including fat, carbohydrates and fiber, while the majority of protein was derived from either plant or animal sources. To our knowledge, this is the first clinical trial to evaluate the impact of protein source on GI health and the gut luminal environment in human subjects. Because this trial was conducted in healthy individuals, it is not known whether the beneficial effects of PP, and conversely the harmful effects of EWP, would influence the pathogenesis of GI disease, such as IBD. Moreover, it is unclear whether PP simply maintains homeostasis, and as such, excludes a potential exogenous factor (*i.e.*, high levels of diet-derived peptides) that could impair GI function or whether its consumption actively protects the gut. It should be noted that exclusive enteral nutrition, which provides protein (and other dietary components) in an easily digestible/absorbable form, can induce remission in patients with IBD (*45–48*). This suggests that limiting dietary protein deposition into the colon could be one of the mechanisms by which this type of diet is beneficial, although its long-term use may have unintended harmful consequences, given that chronic elimination of all dietary peptides impairs immunogenic tolerance to dietary antigens (*49*). Therefore, it stands to reason that consumption of a whole protein source that is digestible (*e.g.*, PP) would maintain homeostatic levels of dietary peptides in the colon while also imparting immunological tolerance. Certainly, the findings shown here, along with recent preclinical work showing that a PP-based diet attenuates colitis in mice (*25*), provide a foundation for directly testing whether using PP to replace other protein sources in diet lessens the severity of IBD or other GI diseases.

## Materials and Methods

### Eligibility criteria

Inclusion criteria included adults 18-30 years of age and a BMI between 18-29.9kg/m², while exclusion criteria consisted of a self-reported history of diabetes, renal, liver, cardiovascular disease, malnutrition, gastrointestinal diseases, including irritable bowel syndrome, inflammatory bowel diseases, chronic constipation or diarrhea, and mental illness, including depression, anxiety, bipolar disorder. In addition, those adhering to vegan or vegetarian diets, having used antibiotics or probiotics within one month of the initiation of the study, were pregnant or planning to become pregnant were excluded.

### Subject Recruitment and Enrollment

Subjects were recruited by email blasts to students and employees of Stony Brook University, Stony Brook, NY. Additionally, recruitment flyers were distributed on campus. These email blasts and flyers contained a link to the survey or QR code directing potential subjects to an initial Qualtricsᵡᴹ (Qualtrics, Provo, UT) survey to determine initial eligibility, willingness to avoid alcohol and commit to only consuming food provided by the study for two, 10-day periods. Potential subjects meeting these criteria were scheduled for an in-person evaluation. After a comprehensive review of the study requirements, each subject was consented, their height and weight measured, and a health survey was completed to confirm eligibility. After providing written informed consent, subjects completed an indirect calorimetry test to determine energy needs (Med Gem® Microlife Calorimeter, Medical Holm Solutions, Golden, CO) as well as instruction on how to complete a three-day food record using the Qualtricsᵡᴹ survey. Subjects were asked to review the study diet menu (**Fig. S7**) and ingredient list and trial sample smoothies for tolerability as part of the consent process. Subjects were given a $400 stipend divided into two payments, received upon completion of each 10-day diet intervention period.

Subjects were weighed three times during each diet intervention period using a balance scale. A total of 261 individuals completed the initial interest surveys and 57 completed the in-person screening, leading to enrollment of 28 subjects, of whom, 18 began the interventions with one subject leaving after the first 10-day stage (**Fig. 1**). The demographic information of the 18 subjects is shown in Table S3. This study was approved by the Institutional Review Board at Stony Brook University (SBU) (IRB# 2022-00427) and registered on Clinicaltrials.gov prior to recruitment of subjects (NCT05619939).

### Human diet formulation and provision

The nutrient content of the subjects’ typical diet, based on the 3-day Qualtrics food/beverage log was analyzed using Food Processor® software. These results and indirect calorimetry were used to inform on the composition of study meals for each subject to ensure adequate nutrition. Meals were prepared by a professional chef and registered dietitian in a metabolic kitchen housed in SBU’s Business Incubator in Calverton, NY. Subjects were provided with individualized very low protein diets (VLPD) supplemented with smoothies containing pea or egg white-derived protein isolates (Judees©). The VLPD consisted of starches, fruits, vegetables and fats, and provided approximately 30% of total protein, while the remaining 70% of protein was provided through the smoothies (**Fig. S7**). Protein shakes were consumed by subjects twice per day to align with breakfast and dinner meals. The combined intake of the VLPD and protein shake was standardized for each subject to provide 1.5g of protein per kg of body weight (Table S1). Subjects were randomized to receive either 10 days of pea protein isolate followed by 10 days of egg white protein isolate, or the same isolates in the opposite order (**Fig. 2**). A two-week washout period was implemented in between provision of each type of isolate (**Fig. 2**). Each day of the feeding period, subjects were provided previously frozen meals for that day in a cooler bag along with snacks, which included directions for safe storage and reheating. Breakfast meals were consumed on site and cooler bags were returned and refilled for the day.

### Assessment of diet adherence

Subjects were instructed to take a picture of each meal before and after eating and upload the pictures *via* a Qualtrics-based survey, in order to estimate how much food was consumed each day, as previously described (*50*). Although subjects were instructed to only eat the food and beverages provided for the study, they logged any consumption of food or beverages not provided by the study (*i.e.*, ‘cheats’), as well as uploaded pictures before and after consuming these non-study foods *via* this same Qualtrics-based survey. Cheats were analyzed by Food Processor to estimate nutrient/calorie content.

### Sample collection and GI health assays

Within 3 days prior to and after each 10-day dietary intervention period, stool samples were collected, lactulose:mannitol tests were performed and GI-PROMIS questionnaires were completed. Stool was collected by subjects using a stool collection kit. If samples were generated during the day, samples were delivered to the research team the same day. If stool was generated in the evening, it was stored in the subjects’ refrigerator overnight and delivered to the research team the following morning. Stool samples were aliquoted into tubes and stored at −80 °C until analysis. The lactulose-mannitol test (Genova Diagnostics) required the subjects to fast overnight and collect a baseline urine sample, from which an aliquot was placed in a collection tube. Subjects then consumed the lactulose/mannitol drink mix and collected all their urine over a six-hour period, from which an aliquot was placed in a separate collection tube, refrigerated, and brought to the research team to be sent for analysis. GI health was evaluated by subject reported assessments through the PROMIS® (Patient-Reported Outcomes Measurement Information System) survey consisting of four domains, including diarrhea (5 items), constipation (9 items), belly pain (6 items), and gas/bloat/flatulence (12 items) (27).

### Metabolomics

Metabolite profiling was carried out at Metabolon, Inc. (Morrisville, NC) using a database of more than 4,500 named molecules, as previously described (*51, 52*). Briefly, fecal samples were lyophilized then resuspended in water (50 μL/mg of dried sample), then homogenized.

Fecal suspensions were used for metabolite extraction using the automated MicroLab STAR system (Hamilton Company, Reno, NV). The resulting extract was divided and used ultra-high performance liquid chromatography-tandem mass spectrometry (UPLC-MS/MS) (positive ionization), UPLCMS/MS (negative ionization), UPLC-MS/MS polar platform (negative ionization). Detection of metabolites was performed using UPLC-MS/MS, as previously described (*53*). Metabolites were identified by automated comparison of the ion features in the experimental samples to a reference library of chemical standard entries that include retention time, molecular weight, preferred adducts, and in-source fragments and MS spectra using software developed at Metabolon, Inc. (*54, 55*). Peaks were quantified using area under the curve and analyzed as described in the statistical analysis section below.

### *In vitro* pepsin-trypsin digestion assay

PP and EWP isolate solutions were diluted into water at 12.5 mg/mL with 35mM NaCl at pH = 2 then incubated with pepsin in a 37°C water bath for 30 minutes. The pH was rapidly raised to 8.4 using NaOH and triethylammonium bicarbonate buffer. Trypsin was then added and the reaction was incubated in a 37°C water bath for 60 minutes. Both enzymes were added at 0.01:1 enzyme:substrate ratios (w/w). Aliquots were collected at sequential time points during pepsin and trypsin incubation, placed into Laemmli buffer and run on a 4-20% gradient SDS-PAGE Tris-Glycine gel. The gel was then fixed for 45 minutes in 12% w/v trichloroacetic acid, stained overnight with 0.1% w/v Coomassie Blue G-250 colloidal stain, and destained. The stained gel was imaged using a BioRad ChemiDoc Imager and densitometry was performed using FIJI (modified NIH ImageJ) software.

### Metagenomics

Metagenomic analysis was carried out at the Microbiome Core Facility of Weill Cornell Medicine as previously described (*56*). Briefly, fecal samples were homogenized in a Qiagen PowerBead 0.1 mm tube using the Promega Maxwell RSC PureFood GMO and Authentication Kit per manufacturer’s instructions. For DNA extraction, 300 µL of homogenate was processed using an automated Promega MaxPrep Liquid Handler instrument loaded with proteinase K, lysis buffer, and elution buffer. DNA was quantified using Quant-iT dsDNA High Sensitivity Assay Kit using a Promega GloMax plate reader. To generate metagenomic shotgun sequencing libraries, the Illumina Nextera XT DNA Library Prep Kit Reference Guide was used.

Library quality & size verification was performed using the Revvity Labchip GX Touch HT instrument with DNA 1K Reagent Kit. Libraries were normalized to 2nM using the Hamilton Microlab NIMBUS instrument. Pooled libraries were sequenced on the Illumina NovaSeq X instrument at loading concentration of 0.13nM + 5% PhiX, paired-end 150bp. Quality-based filtering and removal of host sequences and adapter and quality trimming was performed using kneadData version 0.10.0, using the hg37 human reference genome (https://huttenhower.sph.harvard.edu/kneaddata/). Data were analyzed as described below.

### Metaproteomics

Human and mouse fecal samples were lysed by bead beating in 2 ml Lysing Matrix E tubes (MP Biomedicals). Proteins were extracted from human stool samples by phase separation using a chloroform/methanol followed by filter-aided sample preparation (FASP) (*57, 58*) with some modifications (Supplementary Methods). Proteins were extracted from the mouse fecal samples using the FASP protocol (*58*) with some modifications (Supplementary Methods). The concentration of eluted peptides was determined by a Micro BCA kit (Thermo Scientific, USA) following the manufacturer’s instructions. Peptides were analyzed using 1D-LC-MS/MS analysis performed on a Vanquish Neo UHPLC system (Thermo Scientific) coupled to an Orbitrap Astral (Thermo Scientific) (Supplementary Methods). Proteins were identified using protein sequence databases custom made following the principles described in (*59*). The database constructed in Blakely-Ruiz et al. (*60*) was used to analyze the mass spectra of murine samples. For protein identification, MS/MS spectra were searched against the “human” or “mouse” protein database in Proteome Discoverer v3.3 (Thermo Fisher Scientific).

### Generation and administration of fecal extracts

Fecal pellets were vacuum centrifuged for three hours at room temperature then homogenized by bead beating three times for 20 seconds each in Minimal Essential Medium (MEM) containing 10% dialyzed fetal bovine serum (FBS) and 1% penicillin/streptomycin (P/S) (50 µL per mg dry fecal weight). Samples were centrifuged at 10,000 RCF for 10 minutes at 4°C and supernatant was passed through a 0.22 µm filter. The extracts were stored as single-use aliquots in -80°C until use in assays. IEC-6 cells (ATCC #CRL-1592) (passage 5-12) were seeded at 20,000 cells per well in a 96-well plate using Dulbecco’s Modified Eagle Medium supplemented with 10% heat-inactivated FBS, 1% P/S and 0.1 U/mL human insulin. After 24 hours, cells were treated with complete MEM alone or complete MEM containing fecal extracts for an additional 24 hours (1:200 for human samples and 1:20 for mouse samples).

### 3-[5-dimethylthiazol-2-yl]-2,5 diphenyl tetrazolium bromide (MTT) assay

After incubation with extracts, MTT was added at a final concentration of 5 mg/mL into the existing media and cells were incubated for 4 hours at 37°C. MTT crystals were solubilized with addition of 100 µL per well of 10% sodium dodecyl sulfate in 0.01M HCl, and allowed to dissolve for at least 12 hours at 37°C. Absorbance was measured at 570 nm using the SpectraMax M5 plate reader, and raw values were corrected with a media-only blank samples and made relative to complete MEM alone.

### Protein quantification

Feces were vacuum centrifuged for 3 hours at room temperature. Radioimmunoprecipitation assay buffer was added to the fecal pellet at a concentration of 10 µL/mg of dry feces then homogenized with bead-beating, followed by centrifugation and the supernatant was used for protein quantification. Protein concentration was quantified in both sample types using the BioRad DC Assay per manufacturer instructions. Protein levels detected in MEM + 10% dialyzed FBS were subtracted to calculate the final concentration of fecal extracts.

### Fecal extract size exclusion

Samples were applied to Amicon 3 kDa MWCO Spin Columns, per the manufacturer’s protocol, with an initial sterilization using 70% ethanol. The >3 kDa concentrated retentate was volume-adjusted based on the ratio of the initial extract volume to the volume of concentrated retentate prior to use in experiments. The two fractions were additionally combined to generate a ‘total’ extract that was used as a control group for experiments. Protein quantification was carried out as described above for each fraction to validate efficacy of fractionation.

### Mouse studies

8-week-old C57BL/6J male mice (The Jackson Laboratory) were provided experimental diets *ad libitum* for seven days, following two weeks of bedding swapping to homogenize microbiota. Fecal pellets were collected and flash frozen for subsequent protein quantification and generation of extracts. Mice were then given 1% dextran sodium sulfate *via* drinking water while continuing experimental diets, during which time body weight, diarrhea, and fecal bleeding scores were measured, as previously described (*25*). These studies were approved by the Institutional Animal Care and Use Committee of Stony Brook University (protocol #1418336).

## Data availability statement

The mass spectrometry metaproteomics data have been deposited to the ProteomeXchange Consortium *via* the PRIDE (*61*) partner repository with the dataset identifiers PXD081015 (token: l0rMOOoDkifD) (human data) and PXD081083 (token: YsuFHKDd4s0G) (mouse data).

## Statistical analysis

Gastrointestinal symptom scores and lactulose:mannitol ratios from the human randomized crossover trial were analyzed using relative change from pre- to post-intervention, calculated as (post - pre)/pre. Sequence and period/phase effects were evaluated to assess whether treatment order or intervention period influenced outcome changes. Linear mixed-effects models were then used to estimate treatment-associated relative changes, with participant included as a random effect and treatment, sequence (PP first or EWP first), phase, and gender included as fixed effects. Least-squares mean relative changes with 95% confidence intervals were estimated for each protein source, and treatment contrasts were used to compare relative changes between EWP and PP. Separate models were fit for lactulose:mannitol ratio, total gastrointestinal symptom score, and the diarrhea, constipation, belly pain, and gas/bloating symptom domains. Analyses were performed using SAS 9.4 (SAS Institute Inc., Cary, NC), with two-sided p-value < 0.05 considered statistically significant.

Metabolomic data were analyzed as previously described (*62*). Briefly, metabolomic peak area data were log-transformed, batch-normalized and imputed prior to statistical analysis. Metabolites were classified according to ‘superpathway’ (*e.g*., amino acids, peptides, carbohydrates) and ‘subpathway’ (*e.g*., arginine synthesis) based on available literature and with reference to Metabolon’s internal library of purified standards. Relative abundance (assessed by split-split plot contrast tests) was determined using multiple hypothesis testing.

Statistical and principal component analyses were conducted on the Metabolon Integrated Bioinformatics Platform (Metabolon, Inc.) and MetaboAnalystR 6.0 (*63*). An adjusted P value <0.05 and fold-change >1.5 was considered statistically significant.

For metagenomics, taxonomic composition was calculated using MetaPhlAn4 (*64*).

Abundance of gene families and metabolic pathways were quantified using HUMAnN3 (version 3.6) (*65*). Gene families and pathways were renormalized to relative abundances after removing unmapped/ungrouped counts. Differential abundance analysis was performed with MaAslin2 (*66*) and ANCOM-BC2 (*67*), using a mixed model with subject as random effect and a fixed effect for the primary grouping variable. Diversity analysis was performed with QIIME2(*68*).

Analysis of metaproteomic data was performed by controlling the FDR at 0.05 at the peptide-spectrum match (PSM) and protein levels, while Master Proteins were only retained by strict parsimony rules. Proteins were quantified by spectral counting or area under the curve (AUC), depending on the final use case scenario (Supplementary Methods).

Differences in absorbance by MTT assay, protein concentration, and diarrhea/bleeding scores were determined by unpaired t-test with α = 0.05. Data were graphed with GraphPad Prism 11.0.0 and tables were designed in Microsoft Excel 365.

## Acknowledgements

We acknowledge the biostatistical consultation and support provided by the Biostatistical Consulting Core at the Renaissance School of Medicine, Stony Brook University. All metaproteomics LC-MS/MS measurements were made in the Molecular Education, Technology, and Research Innovation Center (METRIC) at North Carolina State University.

## Funding

This work was supported by the Pulse Crop Health Initiative of the US. Department of Agriculture (58-3060-2-034) (D.C.M., J.C.S., O.A.), the National Institutes of Health under Award Numbers R35GM138362 (M.K.) and R01DK118024 (M.K.), and startup funds from the Stony Brook Cancer Center and Bahl Center for Metabolomics and Imaging (D.C.M.).

## Competing Interests

Authors declare no competing interests.

---

**Fig. S1.**
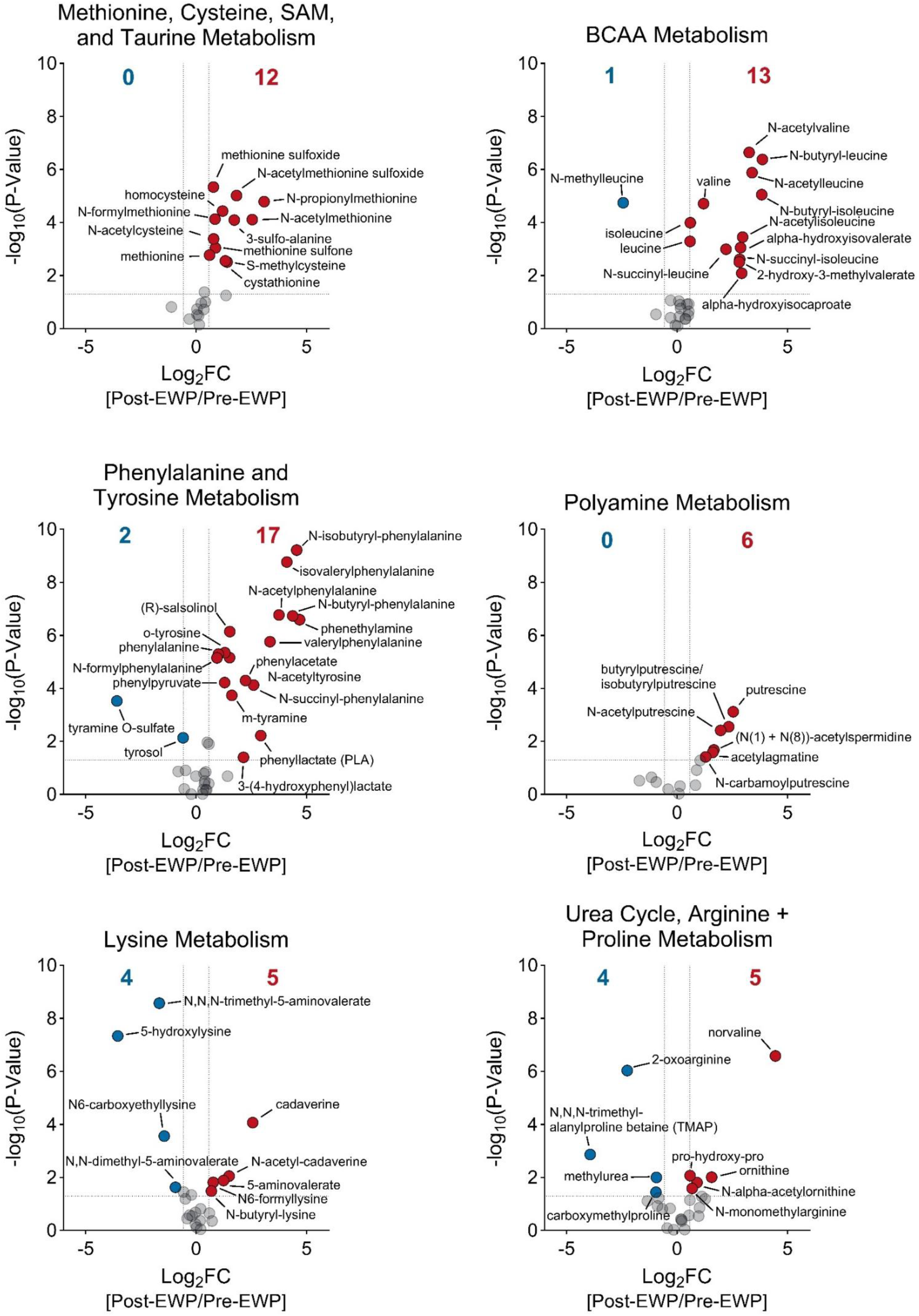
**Egg white protein consumption alters the abundance of metabolites within amino acid-related pathways**. Data generated from metabolomic profiling of feces from human subjects following egg white protein consumption was analyzed by pathway enrichment and the top 6 sub-pathways within the ‘Amino Acids’ superpathway are shown.

**Fig. S2.**
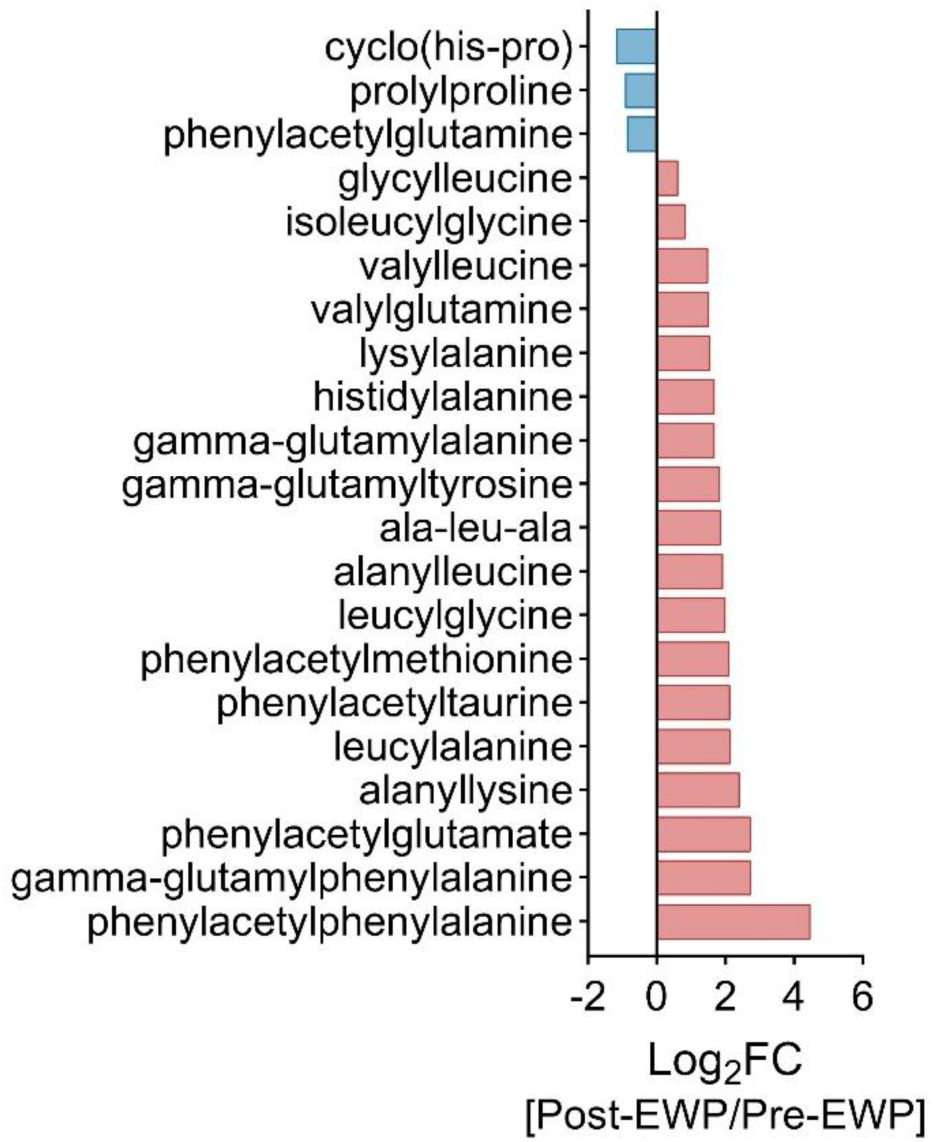
Consumption of an egg white protein-based diet alters the abundance of fecal peptides. Log2 fold-change (FC) of di- and tri-peptide levels in feces of human subjects following 10 days of egg white protein consumption is shown.

**Fig. S3.**
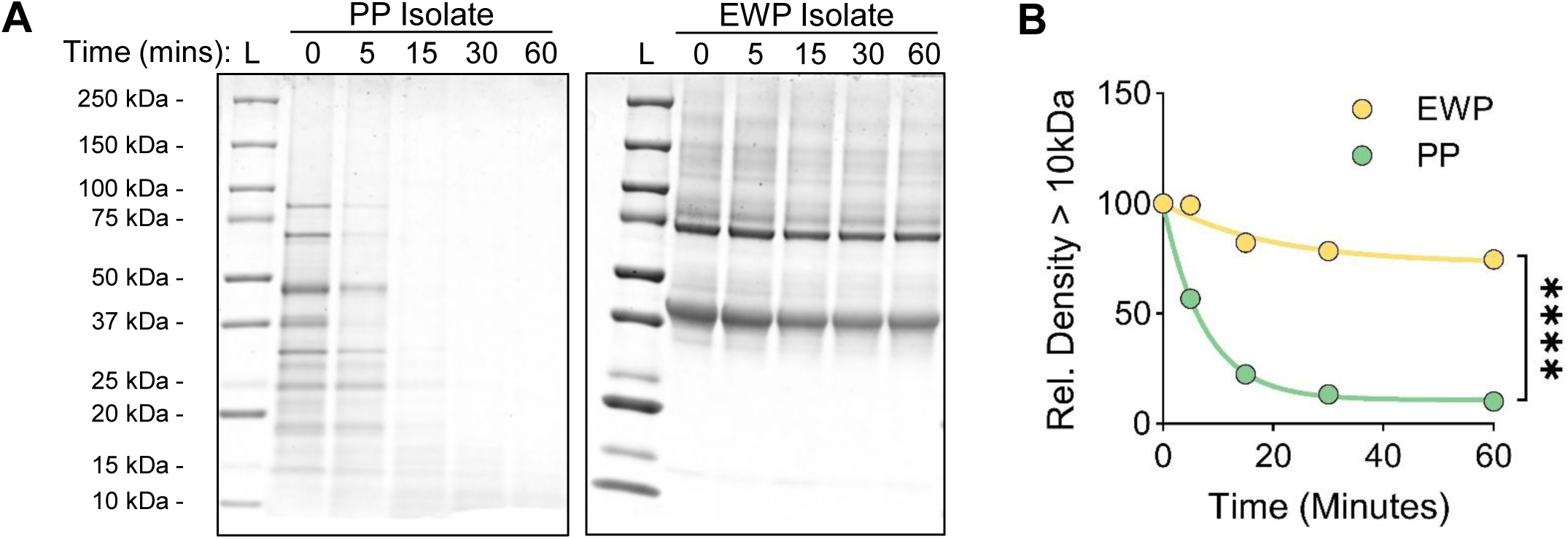
Egg white and pea protein isolates are differentially susceptible to trypsin digestion. Protein isolates from peas (PP) and egg whites (EWP) were incubated with trypsin and the reaction mix was sampled at the indicated time points. Samples were analyzed with SDS-PAGE stained with Coomassie Blue G250 (A) and densitometry was performed using NIH ImageJ (B). ****P<0.0001 determined by a one-phase exponential decay regression test.

**Fig. S4.**
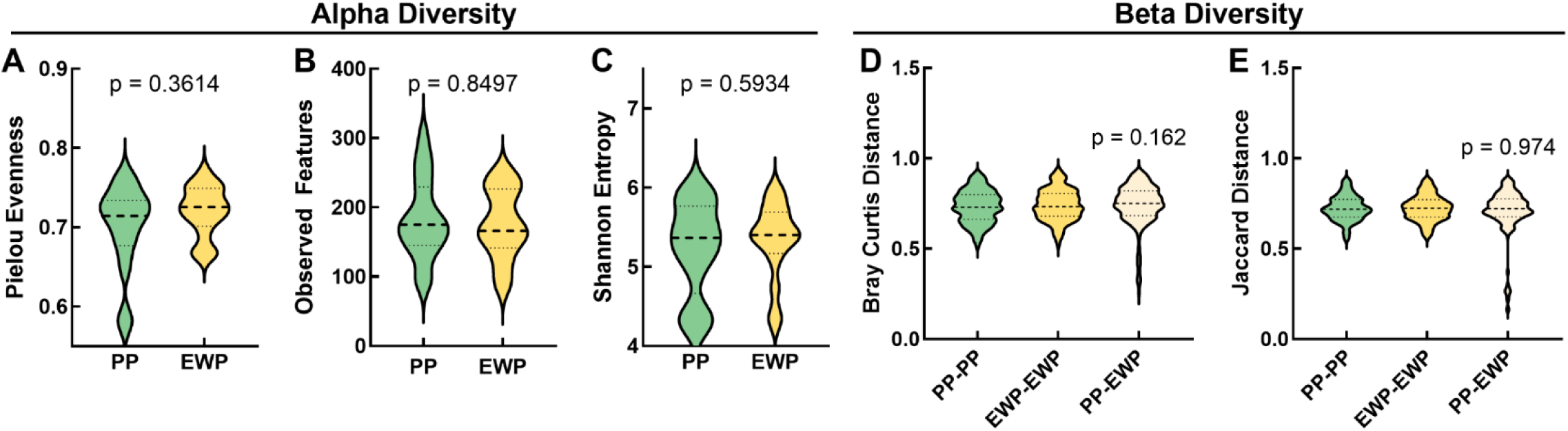
Altering dietary protein source does not impact bacterial diversity. Metagenomic analysis was performed on stool samples collected from subjects consuming diets in which the majority of protein content was comprised of isolates from peas (PP) or egg whites (EWP) and Alpha (A) and Beta (B) diversity were calculated.

**Fig. S5.**
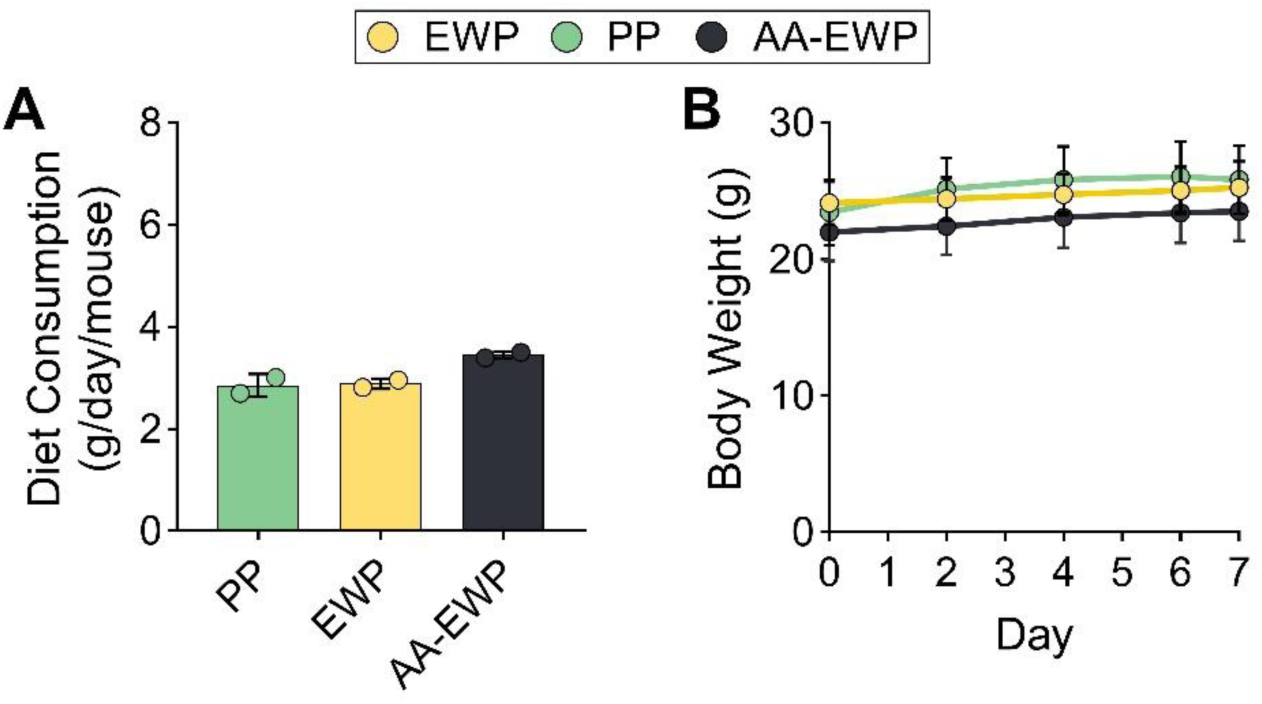
Mice consume comparable amounts of experimental diets and maintain body weight. Male C57BL/6J mice were fed isocaloric, isonitrogenous diets with protein comprised of isolate from peas (PP) or egg whites (EWP) or an amino acid (AA) mix reflecting the AA composition of EWP (AA-EWP) for seven days (n = 8/group). Diet consumption (A) and body weight (B) were monitored during the feeding period.

**Fig. S6.**
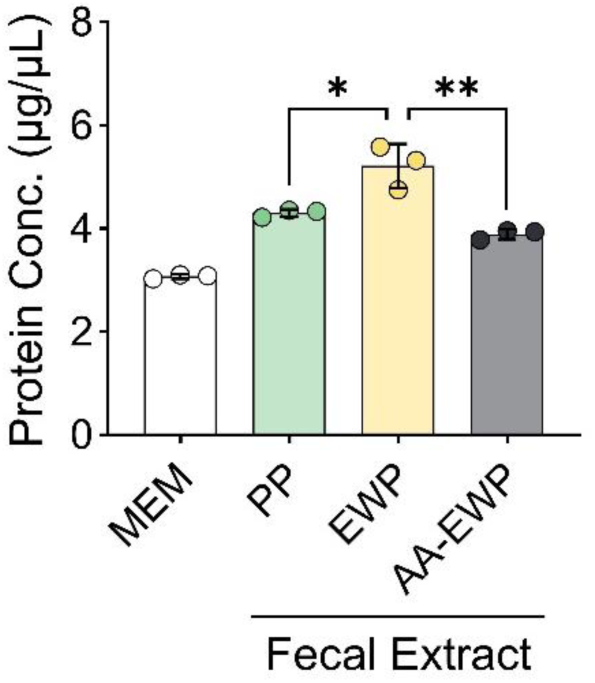
Protein levels vary in fecal extracts from mice fed diets with different protein sources. Protein concentration was measured in fecal extracts (FEs) of samples from mice fed diets containing protein from peas (PP) or egg whites (EWP) or an amino acid (AA) mix matching the AA profile of EWP (AA-EWP) for seven days. *P<0.05; **P<0.01 determined by unpaired t-test.

**Fig. S7.**
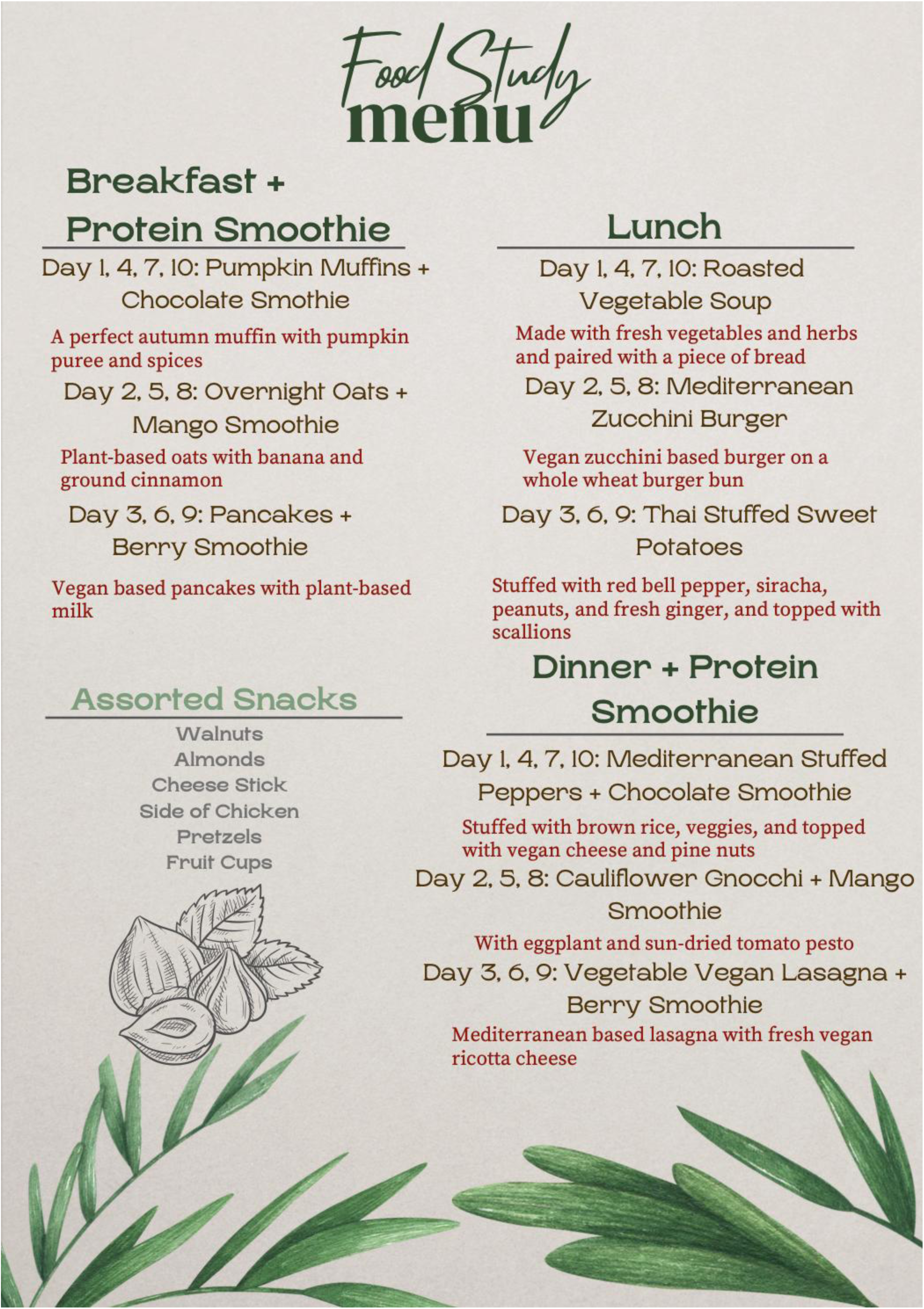
Representative menu of food items used in the trial.

**Table S2.** Composition of murine diets.

| Ingredient (grams) | Pea protein | Egg white protein | Amino acid-based egg white protein |
| --- | --- | --- | --- |
| Pea protein isolate | 209.1 | 0 | 0 |
| Egg white protein isolate | 0 | 210.9 | 0 |
| L-Cystine | 3 | 3 | 8.45 |
| L-Alanine | 0 | 0 | 10.15 |
| L-Arginine | 0 | 0 | 10.86 |
| L-Asparagine-H<sub>2</sub>O | 0 | 0 | 0 |
| L-Aspartate | 0 | 0 | 17.83 |
| L-Glutamic Acid | 0 | 0 | 0 |
| L-Glutamine | 0 | 0 | 22.7 |
| Glycine | 0 | 0 | 6.11 |
| L-Histidine-HCl-H<sub>2</sub>O | 0 | 0 | 4 |
| L-Isoleucine | 0 | 0 | 3.09 |
| L-Leucine | 0 | 0 | 14.3 |
| L-Lysine-HCl | 0 | 0 | 12 |
| L-Methionine | 0 | 0 | 6.67 |
| L-Phenylalanine | 0 | 0 | 10.11 |
| L-Proline | 0 | 0 | 6.07 |
| L-Serine | 0 | 0 | 11.26 |
| L-Threonine | 0 | 0 | 7.76 |
| L-Tryptophan | 0 | 0 | 1.99 |
| L-Tyrosine | 0 | 0 | 6.92 |
| L-Valine | 0 | 0 | 10.8 |
| Corn Starch | 398.5 | 398.5 | 398.5 |
| Maltodextrin 10 | 132 | 132 | 132 |
| Sucrose | 100 | 100 | 100 |
| Cellulose, BW200 | 50 | 50 | 50 |
| Soybean Oil | 51.8 | 70.2 | 71 |
| t-Butylhydroquinone | 0.014 | 0.014 | 0.014 |
| Mineral Mix S10001A (No Ca, P) | 17.5 | 17.5 | 0 |
| Mineral Mix S10020 (No Ca, P, K and Na) | 0 | 0 | 5 |
| Calcium Phosphate, Dibasic | 17.5 | 17.5 | 17.5 |
| Calcium Carbonate | 0 | 0 | 0 |
| Potassium Citrate, Monohydrate | 0 | 0 | 15.7 |
| Potassium Phosphate, Monobasic | 0 | 0 | 0.8 |
| Sodium Bicarbonate | 0 | 0 | 7.5 |
| Sodium Chloride | 19.3 | 12.9 | 17.3 |
| Vitamin Mix V10001 | 10 | 10 | 10 |
| Biotin, 0.1% | 0 | 0.4 | 0 |
| Choline Bitartrate | 2.5 | 2.5 | 2.5 |
| **Total (g)** | **1011.2** | **1016.2** | **1004.81** |
| Protein (g) | 177 | 177 | 177 |
| Carbohydrate (g) | 640.5 | 640.5 | 640.5 |
| Fat (g) | 71 | 71 | 71 |
| Fiber (g) | 50 | 50 | 50 |
| Protein (kcal) | 708 | 708 | 708 |
| Carbohydrate (kcal) | 2562 | 2562 | 2562 |
| Fat (kcal) | 639 | 639 | 639 |
| **Total (kcal)** | **3909** | **3909** | **3909** |
| Protein (g%) | 18 | 17 | 18 |
| Carbohydrate (g%) | 63 | 63 | 64 |
| Fat (g%) | 7 | 7 | 7 |
| Protein (kcal%) | 18 | 18 | 18 |
| Carbohydrate (kcal%) | 66 | 66 | 66 |
| Fat (kcal%) | 16 | 16 | 16 |
| Sodium (g/kg) | 8.8 | 8.8 | 8.8 |
| Calcium (g/kg) | 5.2 | 5.2 | 5.2 |
| Phosphorus (g/kg) | 4.2 | 4.2 | 4.2 |
| Potassium (g/kg) | 5.9 | 5.9 | 5.9 |
| **kcal/gm** | **3.9** | **3.8** | **3.9** |

**Table S3.** Demographics of study subjects.

|  | Number | Percent |
| --- | --- | --- |
| **Sex** |  |  |
| Male | 11 | 64.7 |
| Female | 6 | 35.3 |
| **Ethnicity** |  |  |
| Asian | 9 | 52.9 |
| Black/African American | 2 | 11.8 |
| Caucasian | 3 | 17.6 |
| Hispanic/Latino | 1 | 5.9 |
| Other | 2 | 11.8 |
| Average BMI ± S.D. (kg/m<sup>2</sup>) | 24.7 ± 2.5 | |
| Average Age ± S.D. (years) | 24.2 ± 2.6 | |

## Notes

### Competing Interest Statement

The authors have declared no competing interest.

