## Supplementary Methods for "Diet-derived peptides mediate the effects of dietary protein source on gastrointestinal health"

### Supplemental Methods

**Protein extraction from human samples**

To extract protein from human fecal samples we added 0.5-1 mL of LC-MS grade water to 400-600 mg of sample and homogenized the samples using a hand-held tissue homogenizer (Omni Micro Homogenizer 115 V, Omni International). We extracted proteins by phase separation using a chloroform/methanol extraction followed by filter-aided sample preparation (FASP) [1, 2]. We mixed 100 µL of the homogenized human stool with 1 mL of cold methanol:water 60:40 solution in 2 ml Lysing Matrix E tubes (MP Biomedicals). Samples were bead beat at 6.45 m s^−1^ for 5 cycles of 45 s with 1 minute between cycles. To purify protein, we added 500 µL of cold chloroform to the resulting lysate and placed the samples at -80 °C for 5 min, followed by centrifugation at 4 °C and 12,000 x g for 5 min. The upper and lower layers were removed. We washed the middle protein layer with 500 µL of cold methanol and removed the supernatant using centrifugation. We reconstituted the pelleted protein in 300 µL of 4% (w/v) SDS in 100 mM Tris-HCl, pH 7.6 and stored it at 4 °C overnight. For the FASP peptide preparation, 30 µL of modified SDT buffer (4% (w/v) SDS in 100 mM Tris-HCl pH 7.6, 1.14 M dithiothreitol (DTT)) was added to each protein solution followed by vortexing and heating for 10 min at 95 °C. We centrifuged the samples for 5 min at 21,000 x g to pellet remaining debris. We loaded the protein solutions onto 10 kDa MWCO 500 µl centrifugal filter units (Millipore, Microcon, USA) three times by mixing 60 µL of each protein solution with 400 µL of UA solution (8 M urea in 0.1 M Tris/HCl pH 8.5) followed by centrifugation at 14,000 x g for 20 min. The filters were washed with 200 µL UA, centrifuged at 14,000 x g for 40 min and then incubated for 20 min with 100 µL IAA solution (0.05 M iodoacetamide in UA solution). We removed IAA using a 30 min centrifugation at 14,000 x g. Filters were washed three times with 100 µL UA solution, followed by three washes with 100 µL ABC solution (50 mM Ammonium Bicarbonate), 14,000 x g for 15 min each. Filters were transferred to new collection tubes, and proteins were digested overnight in a wet chamber at 37 °C with 40 µL ABC buffer containing 0.5 µg of MS grade trypsin (Thermo Scientific, USA). The digested proteins were eluted by centrifugation at 14,000 x g for 20 min, followed by the addition of 50 µL of 0.5 M NaCl with 0.4% formic acid, mix at 600 rpm in thermo mixer for 1 min and another centrifugation. The concentration of eluted peptides was determined using a Micro BCA kit (Thermo Scientific, USA) following the manufacturer’s instructions.

**Protein extraction from mouse stool samples**

The mouse stool pellets were lysed by bead beating (5 cycles of 45 s at 6.45 m s^-1^ with 1 min between cycles) in Lysing Matrix E tubes in 550 μL of SDT lysis buffer (4% (w/v) SDS, 100 mM Tris-HCl pH 7.6, 0.1 M DTT). The resulting lysate was heated at 95°C for 10 minutes and centrifuged twice at 21,000 g for 5 minutes to remove the lysing beads and debris. Peptides were then obtained from the lysate using the filter-aided sample preparation method described above, with additional washing steps after the addition of IAA and the subsequent UA washes, in which the filters containing the samples were first washed with 200 μl of ABC buffer, and then washed twice with 200 μl of a 60% Acetonitrile:water solution before the final 3 washes with the ABC buffer and the addition of the trypsin.

**Liquid chromatography and mass spectrometry analysis**

Human and mouse stool peptides were analyzed using 1D-LC-MS/MS analysis performed on a Vanquish Neo UHPLC system (Thermo Scientific) coupled to an Orbitrap Astral (Thermo Scientific). 500 ng of peptide was loaded onto a PepMap™ Neo Trap Cartridge (5 μm particles, 300 μm X 5 mm, Thermo Scientific) and analytical separation was subsequently performed on an EASY-Spray™ PepMap™ RSLC C18 column (2 µm particles, 75 cm x 75 µm ID, Thermo Scientific) using a 70 min gradient with a flow rate of 0.25 ul/min (1%-5% B in 0.1 min, 5%-28% B in 49 min, 28%-40% B in 9 min, 40%-99% B in 2 min, then isocratic at 99% B for 10 min). The eluted peptides were ionized using an Easy spray electrospray source (static spray voltage 2,000 V (+), ion transfer tube temperature 275 °C, radio frequency lens 40%) and analyzed using the Orbitrap Astral mass spectrometer. Mass spectra were acquired in positive ionization mode using data-dependent acquisition (DDA) with the following parameters: full MS1 scans were performed in the Orbitrap with 380-1400 m/z range, 240,000 resolution, 1000% of 1 × 10^7^ AGC target, maximum injection time 200 ms, 1 microscan, profile mode, m/z 445.12003 lock mass. MS/MS scans were performed in the Astral analyzer with 150-2000 m/z range, 80,000 resolution, 50% of 5 × 10^3^ AGC target, maximum injection time 3 ms with 2 microscans, isolation window 1 m/z, normalized collision energy of 32%, cycle time of 1 s, 10 s dynamic exclusion, exclusion of ions of +1 charge state, and centroid mode.

**Database construction**

The protein database for human fecal samples was constructed using three main components: (1) the human proteome (UP000005640, Downloaded July 31, 2025), (2) a custom microbial database and (3) custom diet database. The microbial portion was compiled from the UniRef 90 identifiers quantified using HUMAnN v3 on the metagenomic raw reads sequenced from the same stool samples used for metaproteomics [3]. Briefly the UniRef 90 diamond database for HUMAnN was downloaded using the humann_databases –download function and then the sequences were extracted using the diamond getseq function [4]. We then compiled the sequences for the UniRef90 identifiers quantified in the metagenomes into a microbial protein sequence database and clustered it at 95% ID using cd-hit to remove redundancy [5]. The dietary database was composed of protein sequences of 242 common food species. 335 sequences with 50% similarity with pig trypsin and 1,831 sequences with 50% similarity to 15 most abundant human proteins were removed from the dietary database using cd-hit-2d [6]. Every database (human, microbiota, individual dietary components) was independently clustered with 95% identity threshold using cd-hit [6]. The final database contained 49,042 human, 1,311,254 microbial and 3,759,297 dietary protein sequences. The database was submitted to the PRIDE repository (see section ‘Data availability’). For the mouse stool pellets, the database constructed in Blakely-Ruiz *et al.* was used to analyze the mass spectra [7]. Briefly, the diet-specific mouse databases contained microbial protein sequences from metagenome-assembled genomes constructed by Blakely-Ruiz *et al.*, as well as the reference proteomes of mouse, chicken, and pea from Uniprot.

**Protein identification**

For protein identification MS/MS spectra were searched against the protein database using Proteome Discoverer v3.3 (Thermo Fisher Scientific). Each raw file was searched separately. In the processing step the following nodes were used: Spectrum Files RC with static modification (+57.021 Da) and trypsin as enzyme, Minora Feature Detector, Spectrum Selector, Precursor Detector (1.5 S/N threshold), Top N Peaks Filter (top N 20, Mass Window 100 Da), Sequest HT (Enzyme Trypsin, Precursor Mass 10 ppm, Fragment Mass, 0.05 Da, static modifications carbamidomethylation of cysteine (+57.021 Da), dynamic modifications oxidation of methionine (+15.995 Da), deamidation of asparagine, glutamine, and arginine (+0.984 Da), and N-terminal acetylation (+42.011 Da)), and Percolator for FDR control. We controlled the FDR at 0.05 at the peptide-spectrum match (PSM) and protein levels and only retained Master Proteins by strict parsimony rules. Proteins were quantified by spectral counting or area under the curve (AUC) depending on the final use case scenario.
